# N-acetyl-2-aminofluorene (AAF) processing in adult rat hepatocytes in primary culture occurs by high-affinity low-velocity and low-affinity high-velocity AAF metabolite-forming systems

**DOI:** 10.1101/209072

**Authors:** K.S. Koch, T. Moran, W.T. Shier, H.L. Leffert

## Abstract

N-acetyl-2-aminofluorene (AAF) is a procarcinogen used widely in physiological investigations of chemical hepatocarcinogenesis. Its metabolic pathways have been described extensively, yet little is known about its biochemical processing, growth cycle expression and pharmacological properties inside living hepatocytes ‒ the principal cellular targets of this hepatocarcinogen. In this report, primary monolayer adult rat hepatocyte cultures and high specific-activity [ring G-^3^H]-N-acetyl-2-aminofluorene were used to extend previous observations of metabolic activation of AAF by highly differentiated, proliferation-competent hepatocytes in long-term cultures. AAF metabolism proceeded by zero-order kinetics. Hepatocytes processed significant amounts of procarcinogen (≈12 μg AAF/10^6^ cells/day). Five ring-hydroxylated and one deacylated species of AAF were secreted into the culture media. Extracellular metabolite levels varied during the growth cycle (days 0-13), but their rank quantitative order was time invariant: 5-OH-AAF > 7-OH-AAF > 3-OH-AAF > N-OH-AAF > aminofluorene (AF) > 1-OH-AAF. Lineweaver-Burk analyses revealed two principal classes of metabolism: System I (high-affinity and low-velocity), K_m[APPARENT]_ = 1.64 × 10^−7^ M and V_MAX[APPARENT]_ = 0.1 nmols/10^6^ cells/day; and, System II (low-affinity and high-velocity), K_m[APPARENT]_ = 3.25 × 10^−5^ M and V_MAX[APPARENT]_ = 1000 nmols/10^6^ cells/day. A third system of metabolism of AAF to AF, with K_m[APPARENT]_ and V_MAX[APPARENT]_ constants of 9.6 × 10^−5^ M and 4.7 nmols/10^6^ cells/day, was also observed. Evidence provided in this report and its companion paper suggests selective roles and intracellular locations for System I- and System II-mediated AAF metabolite formation during hepatocarcinogenesis, although some of the molecules and mechanisms responsible for multi-system processing remain to be fully defined.

## INTRODUCTION

Hepatocellular carcinoma (HCC) is the third leading cause of cancer-related deaths worldwide (> 600,000 deaths per year) and the seventh most common cancer (American Cancer Society, 2017; Koch and Leffert, 2015; Mishra *et al.*, 2009; Yang and Roberts, 2010). A wealth of information about the phenomenon of chemical hepatocarcinogenesis has come from investigations of the induction of adult rat HCC (Becker, 1975; Miller, 1975; Weinstein, 1978). Aromatic amines like N-acetyl-2-aminofluorene (AAF) are principal inducers used in these investigations (Kriek, 1974; Weisburger and Weisburger, 1973). Hepatocarcinogenic doses of AAF do not cause tumors when applied topically (Miller *et al.*, 1961). Instead, AAF must first undergo a series of intrahepatic chemical changes. Following rate-limiting N-hydroxylation, sulfate esterification occurs, followed by conversion to more active carcinogenic electrophiles which can form DNA adducts such as dG-C8-AAF and dG-C8-AF (Kriek, 1992; Luo *et al.*, 2000), which ultimately result in HCCs (Bartsch *et al.*, 1972; Cramer *et al.*, 1960; Gutman *et al.*, 1967; Irving, 1962; Kiese *et al.*, 1966; Lotlikar and Luha, 1971; Miller *et al.*, 1960; Morris *et al.*, 1956; Weisburger *et al.*, 1964). The biochemical transformations are performed by hepatocellular enzymes, which are present in the liver at levels higher than in other tissues (DeBaun *et al.*, 1970; Leffert *et al.*, 1977; Santostefano, 1999; Schrenk *et al.*, 1994), particularly by Cyp1A2, which converts AAF to N-OH-AAF. This explains the selectivity ‒ and the utility ‒ of AAF in model studies of rat hepatocarcinogenesis. Other sites, including the bladder, mammary glands and pancreas, are at risk of AAF-induced malignancy, although less frequently (Wilson *et al.*, 1991).

While much is known about the metabolic pathways of AAF, little is known about the cell biology and pharmacology of AAF processing inside living hepatocytes, the major intrahepatic cellular target of this procarcinogen (Koch and Leffert, 2015). A series of experiments were conducted using a well-characterized primary adult rat hepatocyte culture system to quantify the early biochemical processing and biological effects of AAF, and to elucidate the intracellular properties of processing in relation to hepatocyte DNA synthesis and cell division. These investigations are described in this and the subsequent report (Koch *et al.*, submitted, 2017).

This report focuses on AAF uptake and metabolism; the subsequent report (Koch *et al.*, submitted, 2017) focuses on macromolecule binding properties of AAF in relation to hepatocyte proliferation. A third report (Koch *et al.*, in preparation), will describe similar measurements of AAF properties in cultured hepatocytes obtained from livers of adult rats fed a cyclic carcinogenic diet of AAF (Teebor and Becker, 1971).

The results of these studies reveal previously unrecognized aspects of AAF processing, namely the presence of two principal classes of metabolizing systems (high-K_m[APPARENT]_, low-V_MAX[APPARENT]_ and low-K_m[APPARENT]_, high-V_MAX[APPARENT]_) and two principal classes of binding sites (high-K_D[APPARENT]_, Iow-B_MAX[APPARENT]_ and low-K_D[APPARENT]_, high-B_MAX[APPARENT]_). These results have important implications for the mechanisms of AAF processing in relation to hepatocarcinogenesis. Preliminary findings were described elsewhere (Koch and Leffert, 1980; Leffert *et al.*, 1977; Leffert *et al.*, 1983).

## MATERIALS AND METHODS

### Primary hepatocyte culture

Hepatocytes from adult male Fischer 344 rats (180-200g) were isolated by intraportal perfusion with a collagenase cocktail as described previously (Leffert *et al.*, 1977; Leffert *et* al.,1979). Institutional and Animal Care Use Committee regulations at UCSD and NIH guidelines for the care and use of animals were followed. Unless noted, 1 × 10^6^ viable cells (N0) were plated into Falcon^®^ plastic tissue culture dishes (3.5 cm diameter) containing 2 mL arginine-free Dulbecco and Vogt’s Modified Eagle’s Medium. The medium was supplemented with heat-inactivated 15% v/v dialyzed fetal bovine serum, 0.2 mM l-ornithine, and 10 μg each of insulin, inosine and hydrocortisone-succinate/mL. Attached cells (viability > 95%) were recovered by trypsinization, and cell numbers were determined using a Coulter Counter^®^ (Leffert *et al.*, 1979). The reagents and chemicals required for cell culture have been described elsewhere (Leffert *et al.*, 1979). Unless noted, all measurements were made using 3 dishes per point with errors ± 10%.

### Sources and synthesis of radiolabeled AAF

N-acetyl-9-[^14^C]-2-aminofluorene ([^14^C]-AAF) was obtained from New England Nuclear (NEN; Boston, MA [Leffert *et al.*, 1977]). Its purity and specific activity were >98% and 100 dpm pmol^-1^, respectively.

[Ring G-^3^H]-N-acetyl-2-aminofluorene ([^3^H]-AAF) was synthesized from 2-AF (AF or 2-aminofluorene, Aldrich Chemical Co.). The precursor was labeled with tritium by catalytic exchange at NEN. Tritiated precursor (150 mg) was dissolved in 0.3 mL of glacial acetic acid and mixed with 50 mg of platinum catalyst and 25 Ci of tritiated H_2_O. The reaction mixture was stirred overnight at 80°C and the labile tritium removed by evaporation *in vacuo.* The residue was suspended in MeOH, filtered to remove the catalyst and subjected to repeated cycles of dissolution in MeOH and evaporation *in vacuo.* The product of this treatment contained substantial radioactivity which co-migrated with AAF (Aldrich Chemical) as assessed by TLC on silica gel GF plates (Analtech, Inc.) in benzene:acetone [8:2]. The product was saponified by heating an aliquot representing 3 mg starting material for 4 h at 90°C in a mixture of 4 mL EtOH and 8 mL of 50% aqueous KOH. The mixture was cooled and extracted with 40 mL of ether. The ether extract was washed with H_2_O and dried over anhydrous MgSO_4_. The ether was evaporated and [G-^3^H]-2-aminofluorene purified from the residue by preparative thin layer chromatography (TLC) carried out as described above, and using MeOH to elute the material migrating at the same rate as an authentic AAF sample on a parallel track. The MeOH extract was evaporated *in vacuo* and the residue acetylated by dissolving it in 1 mL benzene containing 50 μL of acetic anhydride and allowing it to stand overnight at room temperature. The mixture was evaporated *in vacuo* and [^3^H]-AAF purified from the residue by preparative TLC. The product of this procedure was crystallized from aqueous EtOH. The crystalline material was dissolved in EtOH and stored at −20°C. The specific activity was determined by the optical density at 285 nm compared with a known concentration of an authentic AAF sample. Radioactivity was determined by scintillation counting an aliquot of a dilution in Liquifluor-toluene (NEN). The specific activity of the final [^3^H]-AAF product was 4000 dpm pmol^-1^.

### Isolation and quantification of radiolabeled AAF metabolites

Freshly harvested cell-free culture fluids (2 mL/dish) were incubated with β-glucuronidase and α-amylase, and extracted with acidified organic solvents as previously described (Leffert *et al.*, 1977). Radiolabeled products in the processed extracts (≈65% of the counts) were assayed in parallel by TLC in chloroform:methanol [97:3] and benzene:acetone [4:1], as described previously (Leffert *et al.*, 1977). The two TLC systems were necessary to separate 1 - and 3-position ring-hydroxylated AAF metabolites from parental AAF. Radioactivity in the samples was quantified by scintillation counting as described previously (Leffert *et al.*, 1977). The yields of extracted metabolites that were recovered on TLC plates under both conditions were > 95%. Authentic AAF and AAF-metabolite standards were used for comparison to identify and quantify the co-migrating metabolites in parallel TLC tracks (Leffert *et al.*, 1977).

### Quantification of cellular levels of intracellular free AAF and of covalently bound AAF metabolites

At various times post-plating, [^3^H]-AAF or [^14^C]-AAF were added to the cultures in EtOH vehicle for different intervals of time, as indicated in the Results and Figure Legends. Serial dilutions were made over broad concentration ranges, so that the specific activities of radiolabeled AAF were identical at each AAF concentration. At various times thereafter, the monolayers were washed 3x with 2 mL of ice-cold Ca^2+^- and Mg^2+^-supplemented Tris-HCl buffer, pH.7.4, and extracted with 2 mL of 5% trichloroacetic acid (TCA) overnight in darkness at 4°C. The TCA-soluble extracts were used to quantify the cellular concentrations of free intracellular AAF ([AAF]_i_). The remaining extracts were solubilized (Kruijer *et al.*, 1986), heated at 100°C for 2 min, and brought to room temperature. Covalently bound radiolabeled material in the extracts was precipitated with ice-cold 10% TCA onto Whatman GF/C™ filters. The filters were washed with ice-cold EtOH, air dried, and subjected to scintillation counting to errors of ± 1 % (Leffert *et al.*, 1977). Counting efficiencies of ^14^C and ^3^H were 90-95% and 50-55%, respectively.

### Autoradiography

Twelve day-old cultures were fluid changed into 2 mL fresh complete medium supplemented with 70 μCi of [^3^H]-AAF (2 × 10^−5^ M). Twenty-four hours (h) later, the media were aspirated and the cultures washed 6x with 2 mL Tris-HCl buffer, pH 7.4, and prepared for routine autoradiography by fixation with neutral buffered formalin; by exposure to β-particle sensitive Eastman Kodak™ AR-10 stripping film for 80 days, at 4°C in darkness (Koch and Leffert, 1974), followed by routine development for microscopy.

### Statistical analyses

Linear regression curves and Lineweaver-Burk analyses were calculated using GraphPad™ and Prism™ Software (La Jolla, CA). Mean values, standard deviations (SD) and *P* values were calculated using Microsoft Excel™.

## RESULTS

### Uptake and binding of AAF from the culture medium

AAF was concentrated into cells from the extracellular medium. A log-log plot of the steady-state of intracellular acid-soluble [^14^C]-AAF, [AAF]i, as a function of the initial extracellular concentration of AAF in the medium (6 × 10^−8^ − 2 × 10^−5^ M), [AAF]_o_, yielded a linear absorption curve with a 1^st^ order reaction constant. These results are shown in **Figure 1**.

**Figure 1.**
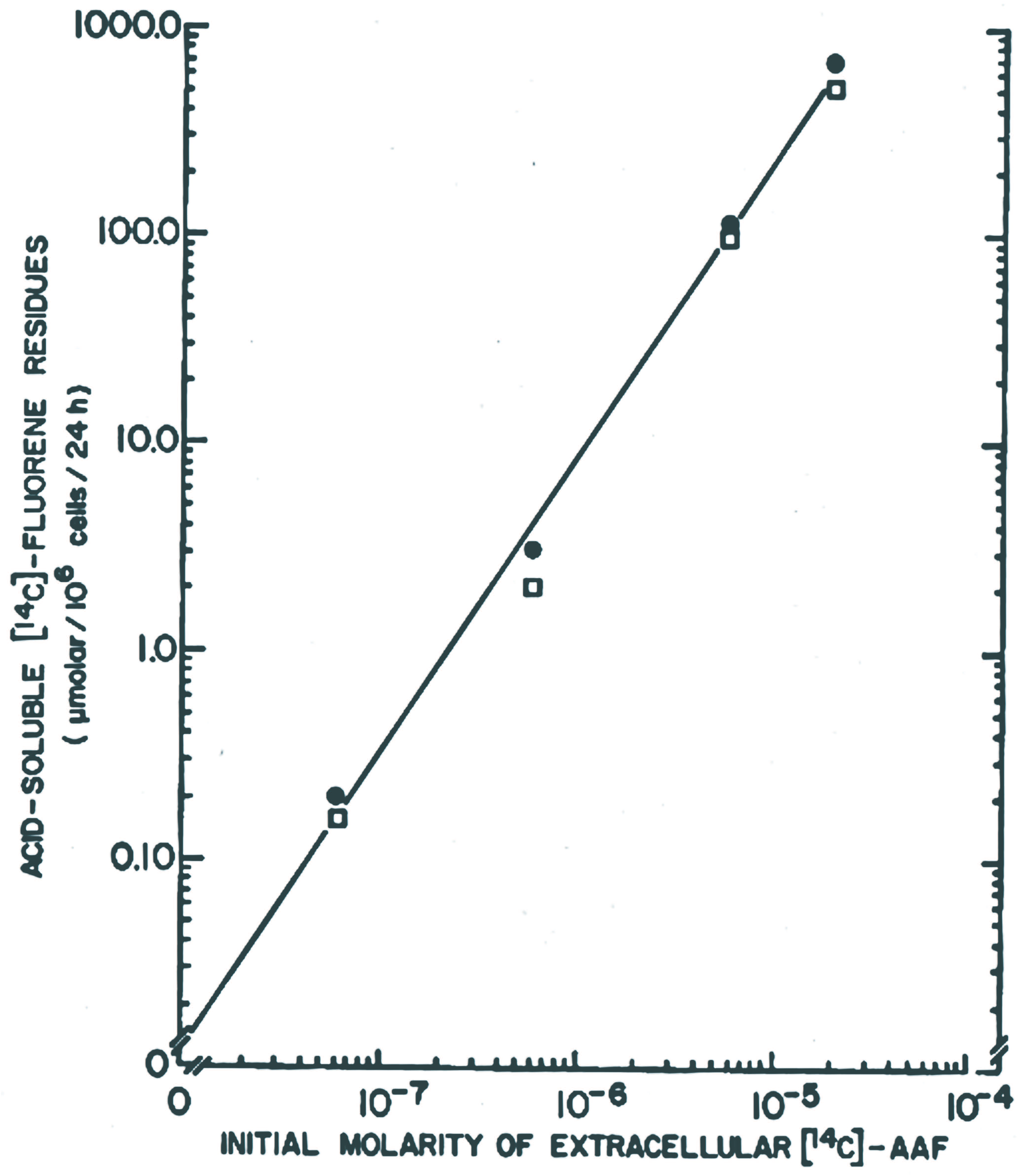
Uptake of AAF from culture media. Acid-soluble [^14^C]-AAF was measured in day 3-4 cultures (N0 = 3.5 × 10^5^ cells/dish) following a 24 h exposure to a range of extracellular AAF concentrations (6 × 10^−8^ M - 2 × 10^−5^ M). Two independent platings were performed (●, □). The results are expressed as soluble fluorene residues (μmolar per 10^6^ cells per 24 h on the y-axis, as a function of the added AAF concentrations on the x-axis. The data were analyzed by linear regression: r = 0.98 (P < 0.0001).

Binding of [^14^C]-AAF to acid-insoluble material was linearly proportional to the numbers of cells per culture (from 0.8 × 10^5^ - 1.0 × 10^6^), and proportional to [AAF]o with a rank order of 2 × 10^−5^ M > 6 × 10^−6^ M > 6 × 10^−7^ M > 6 × 10^−8^ M AAF (**Supplementary Figure 1**). These results indicated that the levels of bound [^14^C]-AAF were not affected by cell density-dependent conditioning of the media (Leffert *et al.*, 1977).

The rates and the levels of binding and uptake of the [^14^C]- or [^3^H]-labeled forms of AAF were virtually identical over an [AAF]_o_ of 6 × 10^−8^ M - 2 × 10^−5^ M. Thus, in 3 day-old cultures, similar quantitative and qualitative results were obtained when either [^14^C]-AAF or [^3^H]-AAF were incubated with the cells over a 24 h period (**Supplementary Figure 2**). Acid-insoluble radiolabeled material in hepatocytes increased 1- to 270-fold in proportion to the [AAF]_o_ (**panel A**, lower and upper curves). By comparison, the levels of acid-soluble [AAF]_i_ were augmented 4- to 1300-fold in proportion to the [AAF]_o_ (**panel B**, lower and upper curves). [AAF]_i_ reached steady states between 6-24 h at all [^14^C]-AAF or [^3^H]-AAF concentrations tested. The specific nature of the bound material and its relationships to [AAF]_i_ are described in the second report (Koch *et al.*, submitted, 2017).

### Kinetics of the disappearance of AAF from the culture medium

When 2 × 10^−5^ M [^3^H]-AAF was added to 3 day-old cultures, 54% of the radiolabeled extracellular AAF disappeared from the culture medium within 24 h (see **inset** to **Figure 2**). The rate of disappearance followed zero-order kinetics (2.3 nmols AAF/10^6^ cells/h ≈12 μg AAF/10^6^ cells/day).

**Figure 2.**
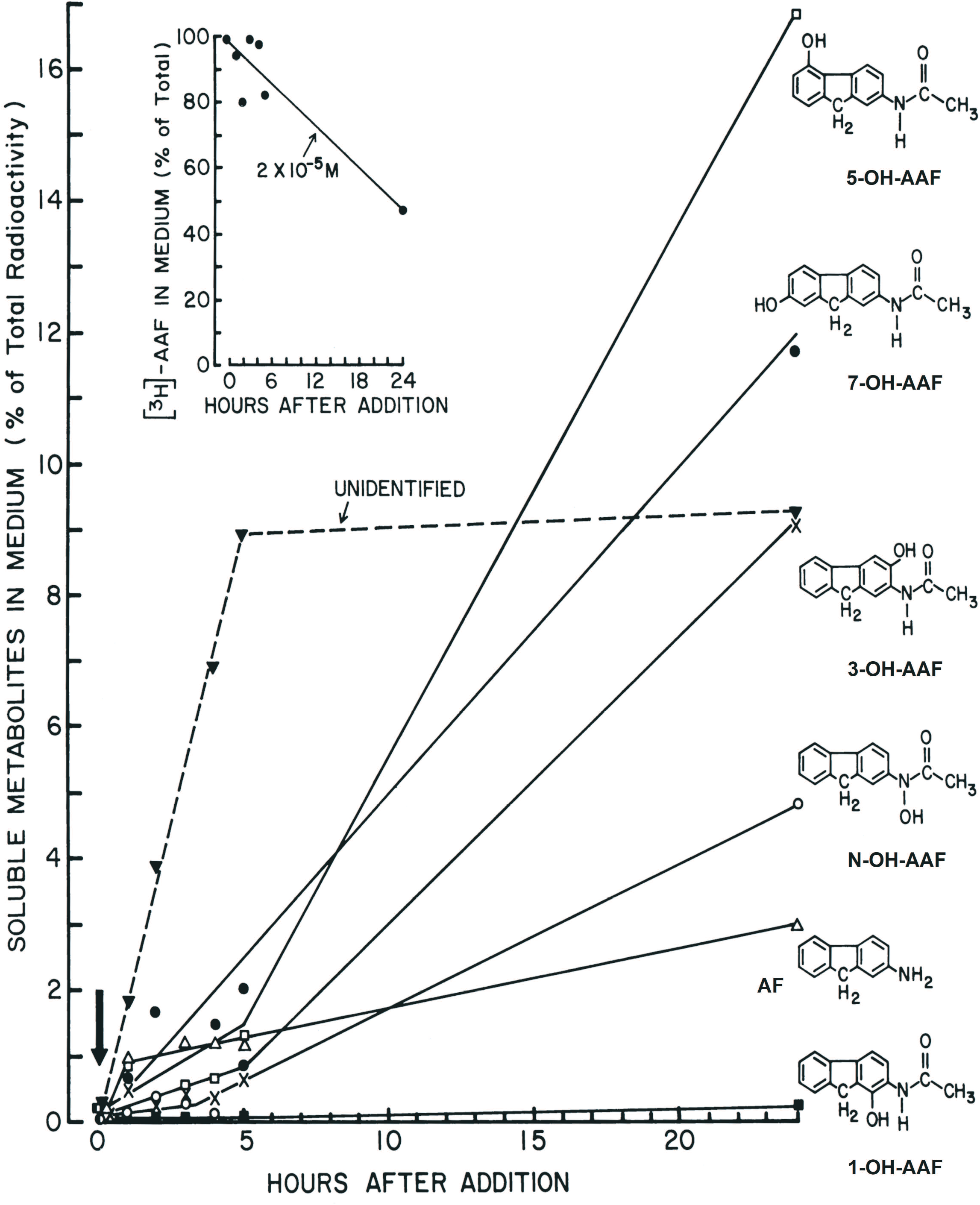
Nature and rates of secretion of AAF metabolites. Three day-old cultures (N0 = 4.0 × 10^5^ cells/dish) were incubated with 40 nmols [^3^H]-AAF (4000 dpm pmol^-1^) per 2 mL medium per dish (initial AAF concentration = 2 × 10^−5^ M). Culture fluids were sampled at the indicated times between zero (↓) at the start of incubation and 24 h (x-axis), and analyzed for AAF disappearance (**inset**), and H_2_O-soluble hydroxylated and deacetylated metabolites (y-axis), as described in Materials and Methods. Each curve is annotated with a contiguous diagram of the structure of each metabolite: □, 5-OH-AAF; ●, 7-OH-AAF; ▾, unidentified at solvent front; **X**, 3-OH-AAF; **O**, N-OH-AAF; Δ, AF; and, ■, 1-OH-AAF (top to bottom). Parameters obtained from linear regression analysis of the **inset** curve: r = 0.92 (P < 0.002).

### Chemical nature and kinetics of secretion of the metabolites of AAF in 3-4 day-old cultures

When 2 × 10^−5^ M [^3^H]-AAF was added to the cultures, six H_2_O-soluble radiolabeled metabolites were secreted into the culture media over 24 h (see the **main panel** of **Figure 2**): **O**, N-hydroxylated-AAF (N-OH-AAF, 4.8% of the total); ●, ring-hydroxylated-AAF (7-OH-AAF), 12%; □, 5-OH-AAF, 16.8%; **X**, 3-OH-AAF, 9.1%; ■, 1-OH-AAF, 0.2%; and Δ, deacetylated aminofluorene (AF, ≈3%). Unidentified metabolites (▼, 9.2%) migrating at the solvent fronts of the TLC plates were also observed. No metabolites of AAF were detected in fresh plating media incubated with [^3^H]-AAF, but without cells, or in centrifuged cell-free conditioned media obtained from unlabeled 3 day-old sister cultures, and subsequently incubated with [^3^H]-AAF for 24 h under regular culture conditions, but without cells. These controls eliminated the possibility that spurious AAF metabolites had formed and were released as a result of chemical oxidation or leakage of P450 enzymes from dying or floating cells in the centrifuged culture media.

Individual metabolites appeared at different rates. At least four were produced as linear functions of time: unidentified ones, from 0-5 h; 7-OH-AAF, from 0-24 h; AF, from 1-24 h, after an initial burst from 0 to 1 h; and 1-OH-AAF from 0-24 h. Within the range of 24 h, only the rate of formation of unidentified metabolites at the solvent front appeared to plateau between 5-24 h. In contrast, no plateaus were observed in the linear rate curves for 7-OH-AAF and 1-OH-AAF within the 24 h period. The metabolites 5-OH-AAF, 3-OH-AAF and N-OH-AAF were secreted biphasically without plateaus. In the latter curves, the low initial rates were linear from ≈ zero-time through 4-5 h. Thereafter, the rates of secretion were augmented 2 to 3-fold at ≈5 h for 5-OH-AAF, 3-OH-AAF and N-OH-AAF, respectively, suggestive of enzyme induction.

### Growth cycle changes in the binding and secretion of metabolites of AAF

Total AAF binding fell exponentially (≈70%) as a function of the culture age during the 0-13 day growth cycle (**Figure 3A**). During the same time interval, this decline was proportional to an exponential decline (≈60%) in the levels of secreted N-OH-AAF (consistent with the rate-limiting role of N-OH-AAF in AAF activation).

**Figure 3.**
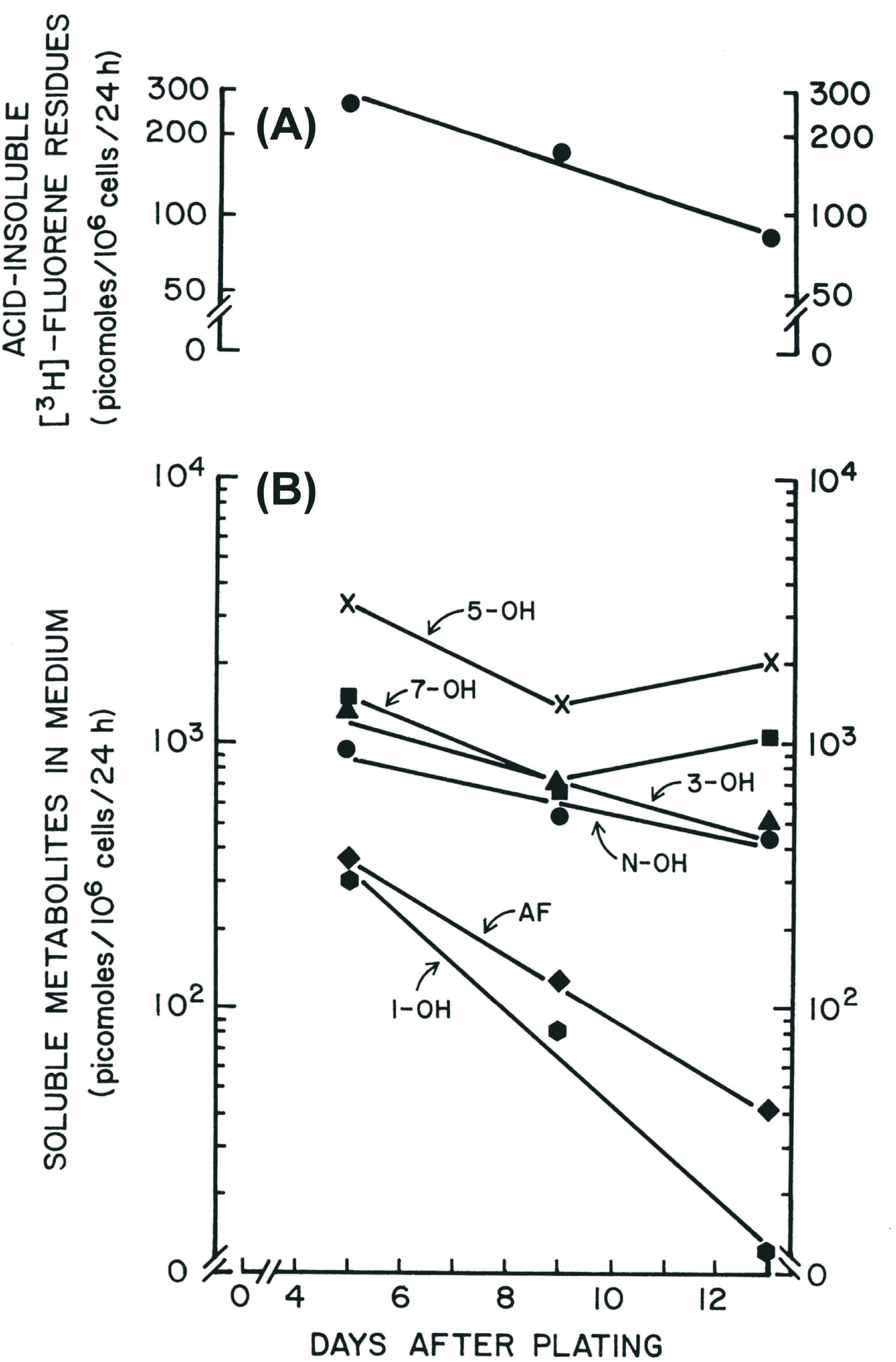
Growth cycle dependence of AAF binding and AAF metabolite secretion. Hepatocytes were plated exactly as described in the **Materials and Methods** subsection, *Primary hepatocyte culture.* On days 4, 8 or 12 post-plating (x-axis [Days After Plating]), the initial plating media were removed and replaced with 2 mL fresh plating media supplemented with 2 × 10^−4^ M [^3^H]-AAF. These culture media were harvested 24 h later on days 5, 9 or 13 per time point (x-axis). **(A)** Acid-insoluble fluorene residues (y-axis, top; picomoles/10^6^ cells/24 h). **(B)** Soluble metabolites in the medium (picomoles/10^6^ cells/24 h), isolated by TLC as described in Materials and Methods (individual metabolites are annotated for each curve; y-axis, bottom). The relative production capacity (%) over the 5-13 day period was determined for each metabolite at each time point from the equation: (100 - [{(% production)_MEASUREMENT DAY_ - (% production)_DAY 5_}/{(% production)_DAY5_}]) × (100).

The overall levels of secreted metabolites varied during the 0-13 day growth cycle, with a rank quantitative order that was unchanged: 5-OH-AAF > 7-OH-AAF > 3-OH-AAF > N-OH-AAF > aminofluorene (AF) > 1-OH-AAF (**Figure 3B**). Between days 5-13, daily production capacities (see the legend to **Figure 3B**) fell exponentially in four instances to 45% (N-OH-AAF), 38% (3-OH-AAF), 11% (AF) and 3% (1-OH-AAF) of the day 5 levels, respectively. In contrast, over the same time interval, 5-OH-AAF and 7- OH-AAF varied biphasically, falling to levels of 48% and 44%, respectively, on day 9, but increasing to levels of 59% and 69%, respectively, by day 13.

### Lineweaver-Burk analyses

The levels of individual metabolites of AAF secreted in 3-4 day-old cultures were proportional to [AAF]_o_ over a 4-log [AAF]_o_ range of 6 × 10^−8^ - 2 × 10^−5^ M. This is shown in **Figure 4**. Two patterns of concentration-dependent curves of metabolite secretion were obtained: AF alone was secreted as a linear curve over the entire concentration range; whereas, 1-, 3-, 5-, 7- and N-OH-AAF followed biphasic secretion curves between 6 × 10^−8^ - 2 × 10^−7^ M and 2 × 10^−7^ - 2 × 10^−5^ M, with the former slope being < the latter.

**Figure 4.**
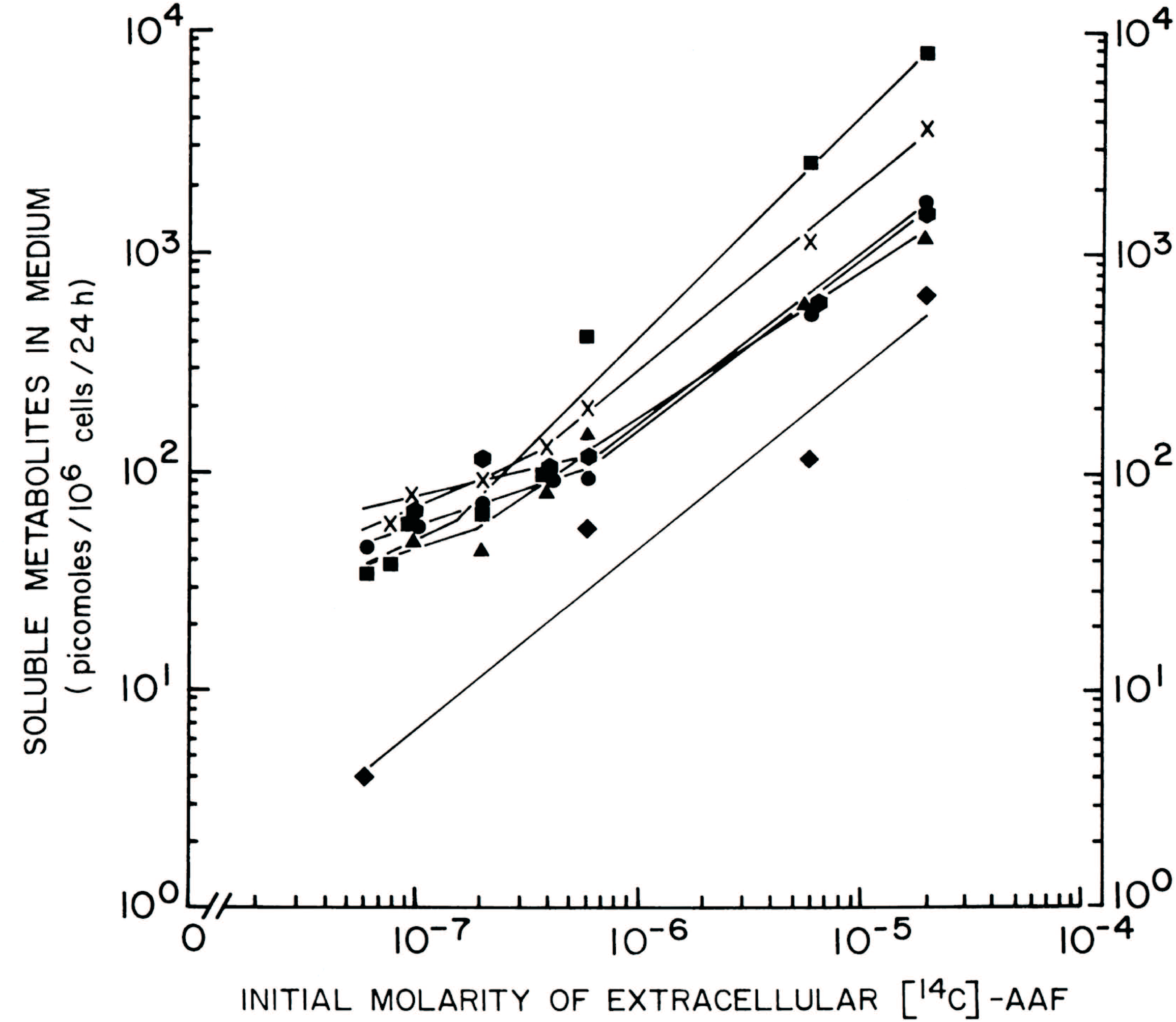
AAF metabolite secretion as a function of the initial AAF concentration. Day 3 cultures (N0 = 3.5 × 10^5^cells/dish) were incubated for a 24 h period with [^14^C]-AAF (6 × 10^−8^ - 2 × 10^−5^ M), at which time the metabolites in the medium were measured. Symbols are the same as those annotated in Figure 5. [S], Concentration of soluble metabolites in the medium (y-axis; picomoles/10^6^ cells/24 h). Initial molarity of extracellular [^14^C]-AAF (x-axis).

When the different data sets were analyzed by Lineweaver-Burk plots (**Figure 5**), the following high-affinity K_m[APPARENT] DAY 4_ constants were obtained: 1.7 × 10^−7^ M, 1.1 × 10^−7^ M, 3.1 × 10^−7^ M, 2.7 × 10^−7^ M, and 1.6 × 10^−7^ M, for metabolites 1-, 3-, 5-, 7- and N- OH-AAF, respectively (collectively, high-affinity K_m[APPARENT] DAY 4_ = 1.64 ± 0.7 (SD) × 10^−7^ M, *P* < 0.05). The following low-affinity K_m[APPARENT] DAY 4_ constants were obtained: 3.9 × 10^−5^ M, 2.6 × 10^−5^ M, 4.4 × 10^−5^ M, 4.0 × 10^−5^ M, and 4.1 × 10^−5^ M for 1 -, 3-, 5-, 7- and N-OH-AAF, respectively (collectively, low-affinity K_m[APPARENT] DAY 4_ = 3.25 ± 0.65 (SD) × 10^−5^ M, *P* < 0.05). Extrapolation of each family of high- and low-affinity K_m[APPARENT] DAY 4_ curves to the y-axis yielded V_MAX[APPARENT] DAY_ 4 values of 0.1 and 1000 nmols/10^6^ cells/24 h, respectively. The curve for the metabolite AF yielded a K_m[APPARENT] DAY 4_ = 9.6 × 10^−5^ M, and a V_MAX[APPARENT] DAY 4_ = 4.7 nmols/10^6^ cells/24 h.

**Figure 5.**
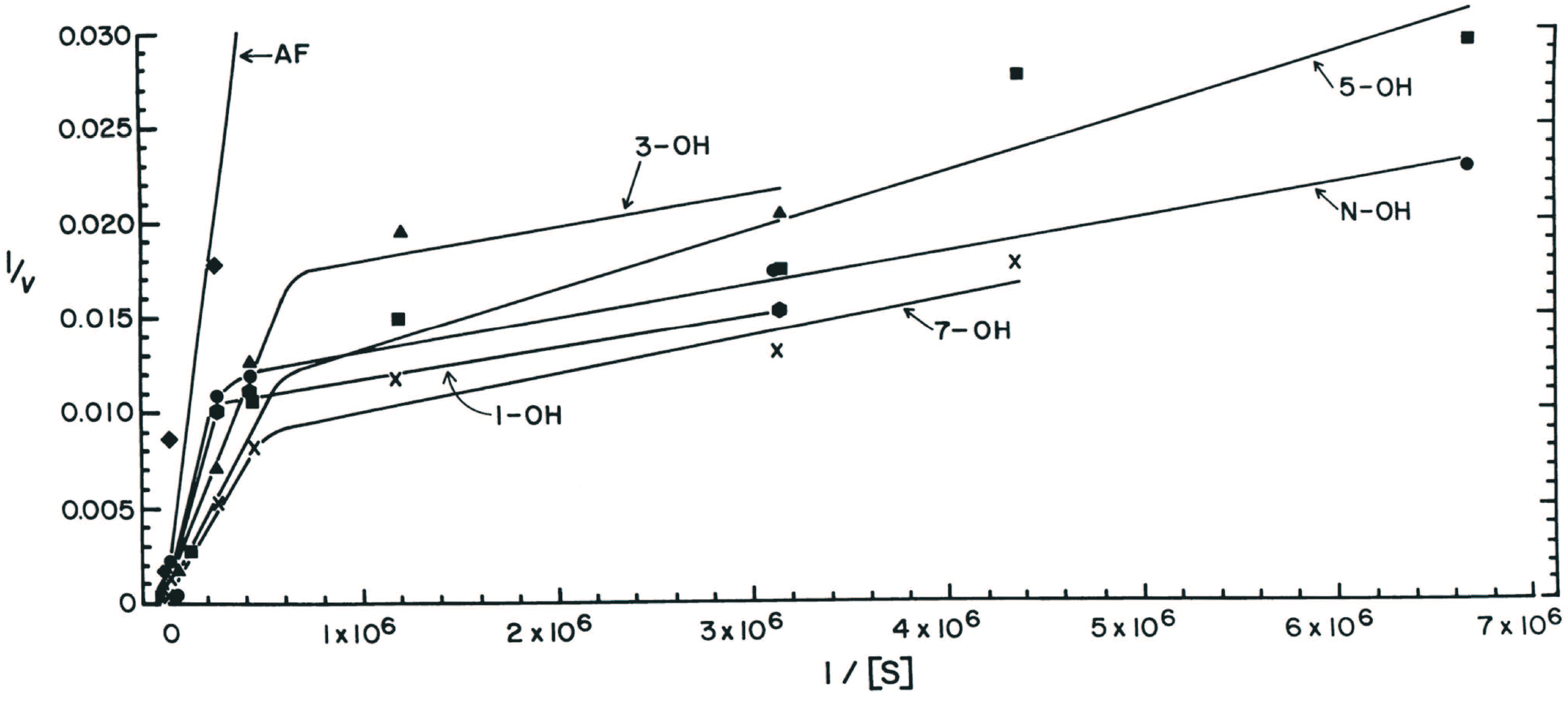
Lineweaver-Burk analyses of AAF metabolite secretion. The 24 h data points were taken from 4 day-old cultures in Figure 4. These data were plotted as: 1/v (y-axis), with v = picomoles/10^6^ cells/24 h; and, 1/[S] (x-axis), with [S] = initial molarity of extracellular [^14^C]-AAF. Metabolites are annotated for each curve in the figure.

### Localization of bound AAF

When quiescent 12 day-old cultures were incubated for 24 h with 2 × 10^−5^ M [^3^H]-AAF and subsequently subjected to autoradiography, the resulting tritium grains (although individually indistinguishable given the length of the development time) localized preferentially to the cytoplasm of hepatocytes in monolayer aggregates (**Figure 6**). Hepatocyte nuclei showed markedly fewer grains than the cytoplasm, consistent with roughly 210-fold reductions in the nuclear levels of covalently bound DNA adducts under these conditions (Koch *et al.*, submitted, 2017); some of these [^3^H]-grains were also detected in mitotic figures. Interspersed non-parenchymal cells showed scant cytoplasmic and far fewer nuclear grains than hepatocytes.

**Figure 6.**
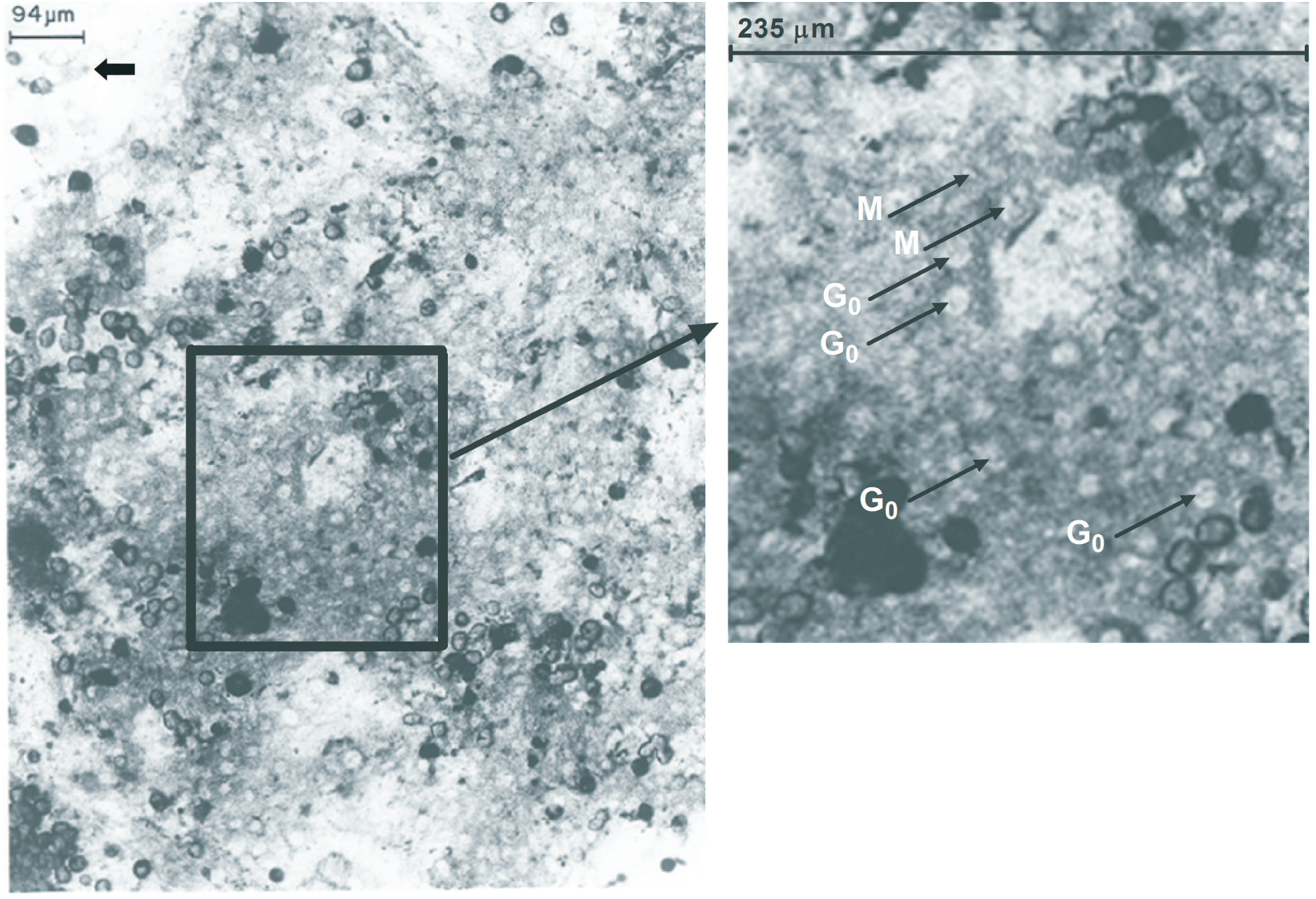
*In situ* autoradiography of covalently-bound metabolites of AAF in day 13 cultures. Hepatocytes were plated exactly as described in the **Materials and Methods** sub-section, *Primary hepatocyte culture.* Twelve days post-plating, the initial plating media were removed and the culture fluids were replaced with 2 mL fresh plating media supplemented with 70 μCi of [^3^H]-AAF (2 × 10^−5^ M). Twenty-four h later, the media were aspirated and the cultures were washed 6x with 2 mL Tris-HCl buffer, pH 7.4, and prepared for autoradiography by fixation with neutral buffered formalin, followed by an 80 day exposure at 4°C in darkness; and subsequent development (Koch and Leffert, 1974). A representative photomicrograph of an unstained culture is shown in the left-hand panel. The short thick arrow (upper left corner) points to a very sparsely labeled non-parenchymal cell. The INSET, enlarged in the right-hand panel, shows [^3^H]-fluorene labeled metaphase (M) chromosomes (long arrows pointing to dark linear structures tilted 45° leftward inside hepatocytes); and G0 cells (long arrows pointing to hepatocytes with densely labeled cytoplasm, and to nuclei with reduced levels of centrally located grains). The **horizontal bars** at the top of each panel give the dimensions in microns along the surface of the monolayer. (Modified from Koch and Leffert, 1980, and used by permission of John Wiley and Sons.)

After 24 h exposure to 2 × 10^−5^ M AAF, cultured hepatocytes (day 13) showed no significant cytotoxicity as revealed by the: a] absence of detached floating cells (Howard *et al.*, 1981); b] levels of DNA replication that were statistically similar to control levels (Koch *et al.*, submitted, 2017), and presence of mitotic hepatocytes (**Figure 6**); and, c] significant conversion of extracellular levels of 2 × 10^−4^ M AAF to the soluble metabolite N-OH-AAF (**Figure 3**), and of radiolabeled AAF to covalently bound macromolecular DNA and protein adducts (Koch *et al.*, submitted, 2017).

## DISCUSSION

The investigations reported here have revealed several aspects of the processing and pharmacological properties of the formation of AAF metabolites, key physicochemical constants, and the two principal locations of AAF metabolism inside normal primary adult rat hepatocytes in long-term cultures. Metabolite formation was surveyed over an [AAF]_o_ range of 2 × 10^−8^ M - 2 × 10^−4^ M, wider than the concentration ranges reported previously (Guzelian *et al.*, 1977; Michalopoulos and Pitot, 1975; Michalopoulos *et al.*, 1976; Monteith *et al.*, 1988; Spilman and Byard, 1981).

A 1^st^ order AAF absorption reaction constant suggested that AAF uptake occurred by simple diffusion across plasma membranes as a function of the partition coefficients of the lipid phases. However, as supported by kinetic studies and by Lineweaver-Burk analyses, a component of carrier-mediated transport was also evident as indicated by the appreciable concentration of TCA-soluble [^14^C]-fluorene residues inside cells. Zero-order metabolism of AAF (2 × 10^−5^ M) indicated that enzymatic machinery was present in excess, in agreement with a previous report (Spilman and Byard, 1981).

AAF metabolism into radiolabeled soluble and bound intracellular fluorene residues depended upon cell numbers, growth-state and culture age. These dependencies could not be attributed to the use of the different isotopes of AAF, since [^3^H]-AAF (ring G-^3^H]-N-acetyl-2-aminofluorene) and [^14^C]-AAF (N-acetyl-9-[^14^C]-2-aminofluorene) appeared to be utilized identically by the cells under various conditions (consistent with the chemical location of the bulk of the ^3^H on the C9-position of the fluorene ring). Other metabolites were undetectable and uncharacterized: for instance, 9-OH-AAF, and an ‘unidentified’ group, respectively; HPLC analyses will be needed to detect and identify them (Ryzewski and Malejka-Giganti, 1982). The secretion of N-OH-AAF, the rate limiting step in procarcinogen activation and evidence of sustained hepatocyte differentiation, was observed through day 13; varying results were obtained in other primary hepatocyte systems (Guzelian *et al.*, 1977; Michalopoulos and Pitot, 1975; Michalopoulos *et al.*, 1976; Monteith *et al.*, 1988; Spilman and Byard, 1981). However, the origin of hepatocytes and the conditions of long-term culture used here differed from those of the short-term cultures.

Pharmacologic evidence of two principal AAF metabolizing systems was observed: System I (high-affinity and low-velocity), with a K_m[APPARENT]_ = 1.64 × 10^−7^ M and V_MAX[APPARENT]_ = 0.1 nmols/10^6^ cells/24 h; and System II (low-affinity and high-velocity), with a K_m[APPARENT]_ = 3.25 × 10^−5^ M and V_MAX[APPARENT]_ = 1000 nmols/10^6^ cells/24 h. AF metabolites were generated from AAF by a third system with a K_m[APPARENT]_ and V_MAX[APPARENT]_ day 4 of 9.6 × 10^−5^ M and 4.7 nmols/10^6^ cells/24 h, respectively.

The biochemical bases of these observations are currently unclear. However, the third metabolism system, which generates AF from AAF, probably involves an arylacetamide deacetylase (AADA) associated with the endoplasmic reticulum of hepatocytes (Trickett *et al.*, 2001). Hepatocyte AADA plays roles in the assembly of very low density lipoprotein (VLDL), an extracellular inhibitor of normal rat hepatocyte proliferation (Leffert and Weinstein, 1976). Serum VLDL levels are inversely correlated with hepatocyte proliferation (Leffert and Weinstein, 1976; Leffert and Koch, 1977) and significant reductions are reported in human patients with HCC (Ooi *et al.*, 2005).

One plausible explanation for the two principal AAF metabolizing systems is that alternatively spliced Cyp1A2 isozymes, like the single-copy human gene (Schweikl *et al.*, 1993; Allorge *et al.*, 2003; Zhou *et al.*, 2010; GeneCards Human Gene Database, 2017), operate in adult rat hepatocytes: one in the nucleus (System I [with one set of K_m_ and V_MAX_ values]); and, the other in the cytoplasm (System II [with another set of K_m_ and V_MAX_ values]). Although neither AAF metabolism via non-parenchymal cells (≤ 5% of the total population) nor bystander effects (Koch and Leffert, 2015) were rigorously excluded, autoradiography with [^3^H]-AAF demonstrated dissimilar distributions of covalently bound intracellular macromolecular adducts (cytoplasm >> nucleus), supporting N-OH-AAF generating Cyp1A2 metabolic activity at both hepatocellular sites. However, little is known about the sub-cellular locations of rat and human hepatocyte Cyp1A2 isozymes (Annalora *et al.*, 2017); therefore the autoradiographic findings are also consistent with one differentially distributed isozyme, wherein the nuclear and cytoplasmic milieu dictate the properties of the K_m_ and V_MAX_ of the enzyme.

In fact, partial nuclear localization of Cyp1A2 is predicted from searches using cNLS Mapper setting cut-off scores at 7.0 (Kosugi *et al.*, 2009). This program identifies two 90% homologous monopartite nuclear localization signals (NLSs [Nigg *et al.*, 1991]) at *aa* positions 451-460 *(glgkrrcige)* and 452-461 *(gmgkrrcige)* in the 75% homologous alignments of the entire rat (GenBank NP_036673.3) and human (UniProt P05117) proteins, respectively; each decapeptide NLS is 53 amino acids from its C-terminus (**Supplementary Figure 3**), far from the enzymes’ active sites (Zhou *et al.*, 2010); and, as expected for NLSs, both have basic isoelectric points of 9.803 (Kozlowski, 2016). Observations in the accompanying report further support intranuclear metabolism of AAF directed by NADPH-dependent Cyp1A2 (Koch *et al.*, submitted, 2017).

Alternatively, the concerted operation of several cytochrome P450s might underlie the observations reported here. For example, apart from the rat Cyp1A2 isozyme, (NP_036673.3), rat hepatic microsomal extracts contain five additional cytochrome P450s (designated Cyp2B1 [NP_001128316.1], Cyp1A1 [NP_036672.2], Cyp2B2 [NP_001185605.1], Cyp2C6 [P05178.2], and Cyp3A2 [NP_695224.2]); and, in cell-free systems reconstituted with purified microsomal proteins, the Cyp1A1 and Cyp1A2 isozymes produce most but not all of the hydroxylated metabolites of AAF (Guengerich *et al*., 1982; Aström *et al.*, 1986; Nebert *et al*., 1991). With the exception of rat Cyp1A2, cNLS Mapper failed to detect NLS domains in the primary sequences of the five additional cytochrome P450s (data not shown). Studies with human liver microsomes have also shown biphasic kinetics for 7-OH-AAF formation, further suggesting that individual metabolites might be produced by different P450s (McManus *et al.*, 1983). Together with enzyme induction of 5-OH-AAF, 3-OH-AAF and N-OH-AAF, mixed activation of different subtypes might have accounted for the U-shaped curves of 7- and 5-OH-AAF metabolite formation during the growth cycle.

It is also possible that differential substrate accessibility, particularly of AAF, N-OH-AAF, and AF, but also potentially of the other metabolites, might occur inside hepatocytes, because altered enzyme activity and polypeptide composition of rat hepatic endoplasmic reticulum have been reported following acute exposure to AAF (Kaderbhai *et al.*, 1982). Further investigations will be required to address these unresolved issues.

## ACKNOWLEDGMENTS

We thank Hal Skelly for additional technical assistance.

---

**Supplementary Figure 1. Cell density dependence of [^14^C]-AAF binding over a 4- log concentration range.** Binding of [^14^C]-fluorene residues (Acid-insoluble radioactivity) was measured after a 24 h exposure between days 3-4, over a [^14^C]-AAF concentration range of 6 × 10^−8^ - 2 × 10^−5^ M (y-axis), as a function of the day 4 numbers of cells per dish (x-axis). Each curve is annotated with the initial concentration of [^14^C]-AAF.

**Supplementary Figure 2. Kinetics of [^14^C]-AAF and [^3^H]-AAF binding and uptake in 3-4 day-old cultures over a 4-log AAF concentration range.** The arrows indicate the time of addition of each isotopic label of AAF over a concentration range of 6 × 10^−8^ M - 2 × 10^−5^ M. Initial concentrations and isotopes of AAF are annotated in the figure ([^3^H]-AAF, open symbols; [^14^C]-AAF, filled symbols). Over a 24 h period (x-axis), measurements were made of **(A)** Acid-insoluble fluorene residues (picomoles per 10^6^ cells); and **(B)** Acid-soluble fluorene residues (μmolar per 10^6^ cells) as expressed on the y-axes (top and bottom).

**Supplementary Figure 3. Evidence of partial nuclear localization by computer analyses of the amino acid sequences of rat and human hepatocyte Cyp1A2.** Rat and human primary amino acid sequences were obtained from Genbank and UniProt databases (<u>https://www.ncbi.nlm.nih.gov/protein?cmd=retrieve</u> and <u>www.uniprot.org/uniprot/P05177</u>, respectively). The two sequences were aligned by standard procedures (<u>https://www.ncbi.nlm.nih.gov/tools/cobalt/re_cobalt.cgi</u>); a consensus sequence lies in between them. Searches for NLS-like domains were performed as directed by cNLS Mapper (http://nls-mapper.iab.keio.ac.jp/cgi-bin/NLSMapperform.cgi) using the entire region and cut-off scores = 7.0. Decapeptide tracts are labeled in the alignments in <u>**bold underlined**</u> text. NLS-like tracts were not identified in the amino acid sequences of five other rat cytochrome P450s (see **DISCUSSION**).

## References

Allorge, D., Chevalier, D., Lo-Guidice, J.M., Cauffiez, C., Suard, F., Baumann, P., Eap, C.B., and Broly, F. Identification of a novel splice-site mutation in the CYP1A2 gene. (2003). Br J Clin Pharmacol. 56, 341–344.

American Cancer Society (2017). <u>http://www.cancer.org/acs/groups/cid/documents/webcontent/003114-pdf.pdf</u>

Annalora, A.J., Marcus, C.B., and Iversen, P.L. (2017). Alternative splicing in the cytochrome P450 superfamily expands protein diversity to augment gene function and redirect human drug metabolism. Drug Metab Dispos. 45, 375–389.

Aström, A., Birberg, W., Pilotti, A., and DePierre, J.W. (1986). Induction of different isozymes of cytochrome P-450 and of microsomal epoxide hydrolase in rat liver by 2-acetylaminofluorene and structurally related compounds. Eur J Biochem. 154, 125–134.

Bartsch H., Dworkin, M., Miller, J.A., and Miller, E.C. (1972). Electrophilic N-acetoxyaminoarenes derived from carcinogenic N-hydroxy-N-acetylaminoarenes by enzymatic deacetylation and transacetylation in liver. Biochim Biophys Acta. 286, 272–298.

Becker, F.F. (1975). In Cancer: A Comprehensive Treatise, Vol. 1, Etiology: Chemical and Physical Carcinogenesis (F.F. Becker, Ed.) Plenum Press, New York and London.

Cramer, J.W., Miller, J.A., and Miller, E.C. (1960). A new metabolic reaction observed in the rat with the carcinogen 2-acetylaminofluorene. J Biol Chem. 235, 885–888.

DeBaun, J.R., Miller, E.C., and Miller, J.A. (1970). N-hydroxy-2-acetylaminofluorene sulfotransferase: its probable role in carcinogenesis and in protein-(methion-S-yl) binding in rat liver. Cancer Res. 30, 577–595.

GenBank (2017). <u>https://www.ncbi.nlm.nih.gov/protein?cmd=retrieve</u>

GeneCards Human Gene Database (2017). <u>http://www.genecards.org/cgi-bin/carddisp.pl?gene=CYP1A2</u>

Guengerich, F.P., Dannan, G.A., Wright, S.T., Martin, M.V., and Kaminsky, L.S. (1982). Purification and characterization of liver microsomal cytochromes P-450: Electrophoretic, spectral, catalytic, and immunochemical properties and inducibility of eight Isozymes Isolated from rats treated with phenobarbital or 1d-naphthoflavone. Biochemistry 21, 6019–6030.

Gutmann, H.R., Galitski, S.B., and Foley, W.A. (1967). The conversion of noncarcinogenic aromatic amides to carcinogenic arylhydroxamic acids by synthetic N-hydroxylation. Cancer Res. 27, 1443–1455.

Guzelian, P. S., Bissell, D.M., and Meyer, U.A. (1977). Drug metabolism in adult rat hepatocytes in primary monolayer culture. Gastroenterology. 72, 1232–1239.

Howard, P.C., Casciano, D.A., Beland, F.A., and Shaddock, Jr., J.G. (1981). The binding of N-hydroxy-2-acetylaminofluorene to DNA and repair of the adducts in primary rat hepatocyte cultures. Carcinogenesis. 2, 97–102.

Irving, C.C. (1962). N-hydroxylation of 2-acetylaminofluorene in the rabbit. Cancer Res. 22, 867–873.

Kaderbhai, M.A., Bradshaw, T.K., and Freedman, R.B. (1982). Alterations in the enzyme activity and polypeptide composition of rat hepatic endoplasmic reticulum during acute exposure to 2-acetylaminofluorene. Chem Biol Interact. 39, 279–299.

Kiese, M., Renner, G., and Weidmann, I. (1966). N-Hydroxylation of 2-aminofluorene in the guinea pig and by guinea pig liver microsomes in vitro. Naunyn Schmiedebergs Arch Exp Pathol Pharmakol. 252, 418–423.

Koch, K., and Leffert, H.L. (1974). Growth control of differentiated fetal rat hepatocytes in primary monolayer culture. VI. Studies with conditioned medium and its functional interactions with serum factors. J Cell Biol., 62, 780–791.

Koch K.S., and Leffert, H.L. (1980). Growth control of differentiated adult rat hepatocytes in primary culture. Ann N Y Acad Sci. 349, 111–127.

Koch, K.S., and Leffert, H.L. (2015). Basic Science of Liver Cancer Stem Cells and Hepatocarcinogenesis, Chapter 14. In Principles of Stem Cell Biology and Cancer, Future Applications and Therapeutics (T. Regad, T. J. Sayers, and R. C. Rees, Eds.), pp. 273–304. John Wiley & Sons, Ltd, Chichester, UK.

Kosugi, S., Hasebe, M., Tomita, M., and Yanagawa, H. (2009). Systematic identification of cell cycle-dependent yeast nucleocytoplasmic shuttling proteins by prediction of composite motifs. Proc Natl Acad Sci U S A. 106, 10171–10176. <u>http://nls-mapper.iab.keio.ac.jp/cgi-bin/NLSMapperform.cgi</u>

Kozlowski, L.P. (2016). IPC - Isoelectric Point Calculator. Biology Direct 11, 55–31 (<u>http://isoelectric.ovh.org/</u>).

Kriek E. (1974). Carcinogenesis by aromatic amines. Biochim Biophys Acta. 355, 177–203.

Kriek, E. (1992). Fifty years of research on N-acetyl-2-aminofluorene, one of the most versatile compounds in experimental cancer research. J Cancer Res Clin Oncol. 118, 481–489.

Kruijer, W., Skelly, H., Botteri, F., van der Putten, H., Barber, J.R., Verma, I.M., and Leffert, H.L. (1986). Proto-oncogene expression in regenerating liver is simulated in cultures of primary adult rat hepatocytes. J Biol Chem. 261, 7929–7933.

Leffert, H.L., and Weinstein, D.B. (1976). Growth control of differentiated fetal rat hepatocytes in primary monolayer culture. IX. Specific inhibition of DNA synthesis initiation by very low density lipoprotein and possible significance to the problem of liver regeneration. J Cell Biol. 70, 20–32.

Leffert, H.L., and Koch, K.S. (1977). Proliferation of hepatocytes. Ciba Found Symp. 55, 61–82.

Leffert, H.L., Koch, K.S., Moran, T., and Williams, M. (1979). [47] Liver Cells. In Methods in Enzymology (S.P. Colowick, and N.O. Kaplan, Eds.). 58, 536–544.

Leffert, H.L., Koch, K.S., Sell, S., Skelly, H., and Shier, W.T. (1983). Biochemistry and biology of N-acetyl-2-aminofluorene in primary cultures of adult rat hepatocytes. In Application of Biological Markers to Carcinogen Testing (H.A. Milman, and S. Sell, Eds.). U S Environmental Protection Agency, Environmental Science Research. 29, 119–133.

Leffert, H.L., Moran, T., Boorstein, R., and Koch, K.S. (1977). Procarcinogen activation and hormonal control of cell proliferation in differentiated primary adult rat liver cell cultures. Nature. 267, 58–61.

Lotlikar, P.D., and Luha, L. (1971). Acetylation of the carcinogen N-hydroxy-2-acetylaminofluorene by acetyl Coenzyme A to form a reactive ester. Mol Pharmacol. 7, 381–388.

Luo, C., Krishnasamy, R., Basu, A.K., and Zou, Y. (2000). Recognition and incision of site-specifically modified C8 guanine adducts formed by 2-aminofluorene, N-acetyl-2-aminofluorene and 1-nitropyrene by UvrABC nuclease. Nucleic Acids Res. 28, 3719–3724.

McManus M.E., Minchin, R.F., Sanderson, N.D., Wirth, P.J., and Thorgeirsson, S.S. (1983). Kinetic evidence for the involvement of multiple forms of human liver cytochrome P-450 in the metabolism of acetylaminofluorene. Carcinogenesis. 4, 693–697.

Michalopoulos, G., and Pitot, H.C. (1975). Primary culture of parenchymal liver cells on collagen membranes. Morphological and biochemical observations. Exp Cell Res. 94, 70–78.

Michalopoulos, G., Sattler, G., Sattler, C., and Pitot, H.C. (1976). Interaction of chemical carcinogens and drug-metabolizing enzymes in primary cultures of hepatic cells from the rat. Amer J Pathol. 85, 755–772.

Miller, E.C. (1978). Some current perspectives on chemical carcinogenesis in humans and experimental animals: Presidential Address. Cancer Res. 38, 1479–1496.

Miller, J.A., Cramer, J.W., and Miller, E.C. (1960). The N- and ring hydroxylation of 2-acetylaminofluorene during carcinogenesis in the rat. Cancer Res. 20, 950–962.

Miller, E.C., Miller, J.A., and Hartmann, H.A. (1961). N-Hydroxy-2-acetylaminofluorene: a metabolite of 2-acetylaminofluorene with increased carcinogenic activity in the rat. Cancer Res. 21, 815–824.

Mishra, L., Mishra, L., Banker, T., Murray, J., Byers, S., Thenappan, A., He, A. R., Shetty, K., Johnson, L., and Reddy, E. P. (2009). Liver stem cells and hepatocellular carcinoma. Hepatology. 49, 318–329.

Monteith, D.K., Michalopoulos, G., and Strom, S.C. (1988). Metabolism of acetylaminofluorene in primary cultures of human hepatocytes: dose-response over a four-log range. Carcinogenesis. 9, 1835–1841.

Morris, H.P., Weisburger, E.K., and Weisburger, J.H. (1956). Urinary metabolites of the carcinogen N-2-fluorenylacetamide. J Natl Cancer Inst. 17, 345–361.

Nebert, D.W., Nelson, D.R., Coon, M.J., Estabrook, R.W., Feyereisen, R., Fujii-Kuriyama, Y., Gonzalez, F.J., Guengerich, F.P., Gunsalus, I.C., Johnson, E.F., Loper, J.C., Sato, R., Waterman, M.R., and Waxman, D.J. (1991). The P450 superfamily: update on new sequences, gene mapping, and recommended nomenclature. DNA Cell Biol. 10, 1–14.

Nigg, E.A., Baeuerle, P.A., and Lührmann, R. (1991). Nuclear import-export: in search of signals and mechanisms. Cell. 66, 15–22.

Ooi, K., Shiraki, K., Sakurai, Y., Morishita, Y., and Nobori, T. (2005). Clinical significance of abnormal lipoprotein patterns in liver diseases. Int J Mol Med. 15, 655–660.

Rat Genome Database (2017). <u>https://rgd.mcw.edu/</u>

Ryzewski, C.N., and Malejka-Giganti, D. (1982). Systems for separation of metabolites of the carcinogen, N-2-fluorenyl-acetamide, by high-performance liquid chromatography. J Chromatogr. 237, 447–456.

Santostefano, M.J., Richardson, V.M., Walker, N.J., Blanton, J., Lindros, K.O., Lucier, G.W., Alcasey, S.K., and Birnbaum, L.S. (1999). Dose-dependent localization of TCDD in isolated centrilobular and periportal hepatocytes. Toxicol Sci. 52, 9–19.

Schrenk, D., Gant, T.W., Michalke, A., Orzechowski, A., Silverman, J.A., Battula, N., and Thorgeirsson, S.S. (1994). Metabolic activation of 2-acetylaminofluorene is required for induction of multidrug resistance gene expression in rat liver cells. Carcinogenesis. 15, 2541–2546.

Schweikl, H, Taylor, J.A., Kitareewan, S., Linko, P., Nagorney, D., and Goldstein, J.A. (1993). Expression of CYP1A1 and CYP1A2 genes in human liver. Pharmacogenetics. 3, 239–249.

Spilman, S.D., and Byard, J.L. (1981). Metabolism of 2-acetylaminofluorene in primary rat hepatocyte cultures. J Toxicol Environ Health. 7, 93–106.

Teebor, G.W., and Becker, F.F. (1971). Regression and persistence of hyperplastic hepatic nodules induced by N-2-Fluorenylacetamide and their relationship to hepatocarcinogenesis. Cancer Res. 31, 1–3.

Trickett, J.I., Patel, D.D., Knight, B.L., Saggerson, E.D., Gibbons, G.F., and Pease, R.J. (2001). Characterization of the rodent genes for arylacetamide deacetylase, a putative microsomal lipase, and evidence for transcriptional regulation. J Biol Chem. 276, 39522–39532.

UniProt Consortium (2014). <u>http://www.uniprot.org/uniprot/P04799</u>

Weinstein, I.B. (1978). Current concepts on mechanisms of chemical carcinogenesis. Bull N Y Acad Med. 54, 366–383.

Weisburger, J.H., Grantham, P.H., Vanhorn, E., Steigbigel, N.H., Rall D.P., and Weisburger, E.K. (1964). Activation and detoxification of N-2-fluorenylacetamide in man. Cancer Res. 24, 475–479.

Weisburger, J.H., and Weisburger, E.K. (1973). Biochemical formation and pharmacological, toxicological, and pathological properties of hydroxylamines and hydroxamic acids. Pharmacol Rev. 25, 1–66.

Wilson, R.H., DeEds, F., and Cox Jr., A.J. (1941). The toxicity and carcinogenic activity of 2-acetaminofluorene. Cancer Res. 1, 595–608.

Yang, J.D., and Roberts, L.R. (2010). Hepatocellular carcinoma: A global view. Nat Rev Gastroenterol Hepatol. 7, 448–458.

Zhou, S.F., Wang, B., Yang, L.P., and Liu, J.P. (2010). Structure, function, regulation and polymorphism and the clinical significance of human cytochrome P450 1A2. Drug Metab Rev. 42, 268–354.

